# Machine learning based prediction of functional capabilities in metagenomically assembled microbial genomes

**DOI:** 10.1101/307157

**Authors:** Fred Farrell, Orkun S. Soyer, Christopher Quince

## Abstract

The increasing popularity of genome resolved meta genomics - the binning of genomes of potentially uncultured organisms direct from the environmental DNA - has resulted in a deluge of draft genomes. There is a pressing need to develop methods to interpret this data. Here, we used machine learning to predict functional and metabolic traits of microbes from their genomes. We collated an extensive database of 84 phenotypic traits associated with 9407 prokaryotic genomes and trained different machine learning models on this data. We found that a lasso logistic regression based on the frequency of gene orthologs had the best combination of functional prediction performance and interpretability. This model was able to classify 65 phenotypic traits with greater than 90

## Introduction

Predicting phenotype from genotype remains one of the major challenges in biology [1,2]. Addressing this challenge is particularly relevant for understanding microbial communities, the study of which had been boosted by an increased ability to extract sequence data directly from communities. Technological improvements in DNA sequencing have led to an explosion in the amount of such data generated. In the context of microbial ecology, large-scale metagenomic studies such as the Human Microbiome Project [3], the Earth Microbiome Project [4] and the Tara Oceans Project [5] have systematically sequenced the microbial communities in a huge variety of environments at great depth. Amplicon sequencing, such as of the 16S rRNA gene, allows detailed study of the taxonomic makeup of these communities, while shotgun metagenomic sequencing allows characterisation of all genes present in an environment. Increasing depth of coverage and improvements in genome binning algorithms for clustering contigs into genomes, in particular the use of differential coverage across different samples [6, 7], are allowing more and more full and partial genomes to be assembled from shotgun metagenomic studies. Many of these genomes are novel and belong to uncultured organisms that are never studied in the laboratory. A recent metagenomic study on aquifer systems [8], for example, reconstructed 2540 separate high-quality, near-complete genomes, and claimed to have discovered an astonishing 47 new phylum-level lineages among them.

Converting this exponentially growing sequence data into functional understanding of microbial communities requires us to determine physiological functions from it [9]. This would allow the inference of key functions in microbial communities, and how these functions change over ecological conditions and with time [9]. In turn, this ability, could allow us to discern ecological adaptations in environmental microbial communities, as well as to achieve functional mechanistic models of stability and function [10].

Efforts to achieve phenotype-genotype mapping from environmental sequence data has so far mostly focussed on phylogenetic assignments using the 16S rRNA gene. This highly conserved gene can provide a phylogenetic assignment at the species (or higher) level, which can then be used to infer general functional traits. While this approach has been commonly used to study ecological distribution of microbial functions e.g. [11–13], its premise of a direct association of function with phylogenetic assignment (i.e. ‘functional coherence of microbial taxa’) is questionable (e.g. [14]). The level of taxonomic coherence of function is not clear even for strains of the same species, where functions can show high variability either due to a few genetic changes or even regulatory changes [2]; [15] discusses this for *E. coli*. It is also a common problem that when certain taxonomic groups in a microbial community are found to show direct associations with certain ecological factors (or health state of an host), these groups are so broad that assigning specific functions to them is hard or impossible [16]. Indeed, a study of specific functional traits across microbial taxa has found that many of these traits are dispersed across the phylogenetic tree [2,17]. Even where a specific functional trait is taxonomically coherent, the phylogenetic approach is limited by our ability to assign taxonomy based on the 16s rRNA gene. The extensive accumulation of metagenomics data indicates that we have sampled only a fraction of microbial diversity, and it is not uncommon for such data to result in many unassigned taxa, as for example by the aquifer study described above which described dozens of apparently novel phyla.

While reconstruction of genomes from metagenomics data reveals the limitation of taxonomic functional assignment, it also opens up a new route to functional assignments. In particular, several methods and bioinformatics pipelines are now emerging that aim to go from raw metagenomics sequence data to binned (i.e. predicted) genomes to functions, e.g. [18]. The functional annotation steps in these tools usually considers specific gene(s) that are known to associate with specific functions or metabolic pathways [18], as identified for example in databases such as the Kyoto Encyclopedia of Genes and Genomes (KEGG) [19], the Pfam database of protein families [20] and the NCBI COG database of orthologous genes [21]. This approach circumvents the problems of taxonomic coherence and 16S rRNA gene based assignment to taxa, yet relies heavily on existing categorization of genes into functions as done in the above databases. While these functional gene groupings are mostly based on accumulated knowledge and experimental data on metabolic pathways, they might miss the full set of genes associated with a given function and do not consider functions that cannot be assigned to a few genes or seemingly well-organised pathways. One route to overcome such limitations is to develop extensive databases of phenotypic traits of microbes without necessarily using a pathway-centric view. These functional assignments could then serve as a source to apply statistical approaches to ‘learn’ genetic drivers of those functions using genomes of associated microbes. Efforts in this direction have recently resulted in the compilation of literature-based assignment of functions in microbes, either covering a large selection of functions and organisms [22,23] or specific ones such as methanogenesis [24]. The FAPROTAX database [22], which we focus on here, is based on an extensive survey of the scientific literature. The aim of creating this database was to allow microbes found from 16S rRNA amplicon sequencing to be assigned into functional and metabolic groups, so that functional variation across environemnts could be studied and compared to taxonomic variation. The authors found that the abundance of functional groups was strongly influenced by environmental conditions in a variety of ocean environments [12]. The bulk of the classifications in the FAPROTAX database come from *Bergey’s Manual of Systematic Bacteriology* [25], and it currently contains 84 phenotypic traits associated with 4600 microbial taxonomic groups.

Here, we use this accumulating data on function-organism associations to develop an algorithmic approach to identify genetic markers, or classifiers, of phenotypic functions. To do this, we combined the FAPROTAX database with genomes downloaded from the NCBI and used machine learning to train statistical models able to infer an organism’s traits from a vector of genes annotated to functional groups. Machine learning techniques, such as logistic regression and support vector machines (SVMs) have previously been used in bioinformatics to infer phenotypic information from gene sequences, for example using SVMs on amino acid k-mer frequencies to predict protein function [26–28]. A recent paper [29] achieved good results in this task with recurrent neural networks, which can classify proteins directly from their sequences without the need for feature extraction.

Here, we use the presence of known gene orthologs, for example from the KEGG database, to train models of the phenotype of whole organisms. This work was inspired by a recent software famework, Traitar [30], which uses SVMs to predict microbial traits based on genomic information in the form of copy numbers of Pfam families. Our work differs from Traitar in the use of the highly detailed FAPROTAX database. Traitar utilised the Global Infectious Disease and Epidemiology Online Network (GIDEON) [31] for its phenotypic annotations, and was therefore biased toward pathogenic traits; we instead focus on traits associated with metabolism and environmental niche. Additionally, we have a significantly larger training set- -genomes from all 9407 unique species having a genome classified as ‘full’ in the NCBI database- -whereas Traitar used 234. The size of the training set is usually expected to have a significant impact on the performance of a machine learning model.

We find that these classifiers perform better than simple taxonomic assignments of function, and reveal both known and new genetic drivers of specific functions. Using these resulting classifiers for over 80 functions, we then analyzed three recent, large-scale metagenomics datasets from three diverse environments: anaerobic digesters, ground water aquifers and the ocean. The classifier-based functional analysis of these datasets revealed significant differences in functional properties between environments and conditions.

## Materials and methods

### Databases and preparation of training data

To train our models, we utilized the combination of the recently-published FAPROTAX database of microbial phenotypes and the NCBI genome database. We downloaded all prokaryotic genomes classified as ‘full’ from the NCBI Genome database. We used the taxonomic information available from NCBI to assign them phenotypes using the script ‘collapse_table.py’ which comes as part of the FAPROTAX database [22]. We then called genes in these genomes using Prodigal [32]. We annotated the resulting inferred coding DNA sequences (CDS) both by aligning against against the KEGG database using Diamond BLASTP [33] and by searching with hmmer3 [34] against Pfam [20]. The result is a matrix of organisms and and their copy numbers of either KEGG orthologs or Pfam domains. We have found that using gene copy number rather than simple presence/absence significantly increases classifier performance. The scripts we used to download and process the genomes are available at https://github.com/chrisquince/GenomeAnalysis.git.

### Statistical modelling

To model the link between genotype, in the form of KEGG ortholog copy numbers or Pfam protein families, and phenotype as represented in the FAPROTAX database, we used a variety of machine learning techniques. These are all different ways of learning the relationship between the features (gene copy numbers) and targets (biological functions) from the training data of 9407 NCBI genomes from unique species. To train the algorithms, we first split this data into a training set (75% of the genomes) and a test set (25%), by random sampling. The algorithms were then trained on the training data, and their performance tested on the unseen test data genomes, to check that the relationships learned are generalisable (i.e. that the algorithms have not ‘overfit’ the training data). Below, we describe the algorithms used.

### Logistic regression

We found logistic regression, a commonly used linear model for classification problems, [35,36] to be an effective approach in our case. We scaled all input features to have mean zero and variance one before performing the regression. Since the number of KEGG orthologs (features) was somewhat larger than the number of training examples, overfitting, whereby the model classifies on features of the training set which are very specific to it, was a serious problem. To alleviate this, we used logistic regression with an *ℓ*1 penalty term, also known as LASSO logistic regression [37], whereby large parameters are penalized in such a way that only a few of the features have a nonzero weight. In detail, the method involves adding a penalty term equal to the *ℓ*1-norm of all of the coefficients of the regressor, thereby penalising nonzero terms, so that the optimization problem becomes:

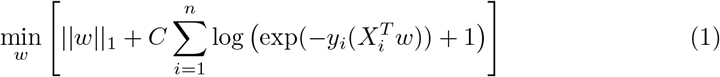

where *w* is the vector of regression weights, *X_i_* are the feature vectors of each example, *y_i_* the classification targets, and *C* is a parameter defining the (inverse) strength of the regularization. This method of regularization is often useful in cases where the number of features is large (similar to or larger than the number of training examples), as most of the features are not used in the classification task. For example, a recent study used *ℓ*1-regularized regression to predict complex human traits such as height and heel-bone density from a large array of SNPs (around 100000), significantly improving on previous estimates of heritability based on individual SNPs [38].

### Random forests

We also used the random forest algorithm, a popular machine learning method which can be applied to both regression and classification problems, which is simple to use, fast and performs fairly well on a wide variety of problems [39]. The random forest is an ‘ensemble’ method, using a collection of slightly randomized classifiers, the results of which are averaged to produce a prediction. This helps to avoid overfitting. A random forest is an ensemble of so-called decision trees. A decision tree is a model which learns to split up training examples into sets according to their feature values, with the aim of separating the target classes. They have the advantage of being invariant under scaling of features and adding of irrelevant features, this last feature being useful in our case where the number of features is very large and many are irrelevant to the classification task; they can also learn more complex relationships between variables than a linear model such as logistic regression. However, an individual tree tends to overfit the training data. A random forest trains a large number of such trees on random subsets of the features and combines these predictions by averaging, avoiding overfitting and much improving performance.

### Support vector machines

Finally, we used support vector machines (SVMs) [?]. An SVM essentially tries to find surfaces in the high-dimensional feature space, which separate the different classes as well as possible, and with as wide a margin as possible between the surface and the examples. These surfaces can be either linear or non-linear (if a non-linear kernel is used); SVMs are therefore capable of learning complex non-linear relationships between features and targets. They can also include regularization terms as in logistic regression, to reduce overfitting.

### Metrics and classifier performance

Since many of the classes which we are attempting to predict are highly unbalanced (e.g. of the 9407 unique species with full genomes in the NCBI database only 83 are hydrogentotrophic methanogens), simple classification accuracy is not a very useful measure of classifier performance. Predicting all labels as negative in the above example would give an accuracy of 99.1% despite not being a useful classifier. We therefore need a metric which can take into account class imbalance. We use the area under the ROC (Receiver Operating Characteristic) curve (AUROC), which is a graph of true postive rate against false positive rate as one varies the cutoff in probability for making a positive prediction [40,41]. An AUROC score much greater than 0.5 (the score for random predictions) indicates a good classifier. In particular, a score of 1 indicates that all positive cases have been assigned a higher probability than all negative cases.

### Prediction of MAG phenotypes

Once classifiers have been trained on the NCBI data, it is possible to use them to make predictions about unseen genomes, such as MAGs generated from shotgun sequencing studies. The MAGs must first be processed to give a matrix of the KEGG ortholog copy numbers associated with them, using the same pipeline as applied above to the NCBI genomes. These matrices are then used as input into the classifiers to produce a matrix of MAGs and their predicted functions, which can be either presence/absence predictions or probabilities.

### MAG collections

We applied our classifier to three separate collections of MAGs from three different studies:

- Tara Oceans MAG collection: This comprised a subset of 660 MAGs from the collection of 957 non-redundant MAGs generated from the Tara Oceans microbiome in Delmont *et al*. [42]. These 660 MAGs were those which had at least 75% of the 36 single-copy prokaryotic core genes identified in Alneberg *et al*. [6] in a single-copy and can thus be considered reasonably complete and pure prokaryotic genomes. The Tara Oceans microbiome survey generated 7.2 terabases of metagenomic data from 243 samples across 68 locations from epipelagic and mesopelagic waters around the globe [43], Delmont *et al*. extracted their MAGs from a subset of 93 of these samples, 61 surface samples and 32 from the deep chlorophyll maximum layer. Therefore these MAGs represent a substantial sample of planktonic microbial life.
- Anaerobic digester (AD) MAG collection: This comprised a collection of 153 MAGs that were constructed by co-assembly and binning of 95 metagenome samples taken from three replicate laboratory anaerobic digestion (AD) bioreactors converting distillery waste into biogas. They were assembled with `Ray` using a kmer size of 41 and all 186,081 contig fragments greater than 2*kbp* in length were clustered by `CONCOCT` [6] generating a total of 355 bins of which 153 were 75% pure and complete and used in this analysis.
- Candidate phyla radiation (CPR) MAG collection: This collection of 581 MAGs is a subset of 797 MAGs provided by the authors of Brown *et al*. 2015 [44]. They comprise members of the Candidate phyla radiation (CPR) assembled from ground water enriched with acetate.

## Results

To address functional prediction of meta genome assembled genomes (MAGs), we aimed to develop here a machine learning approach based on known phenotypic functions and genomes harbouring those. We collated a database of genomes and their known functions (see Methods), and used this to train different machine learning approaches for prediction (see Methods).

### Classification accuracy

Figure 1 shows the performance of the different machine learning algorithms in the classification task on the test set in terms of AUROC (area under the Receiver Operating Characteristic, see Materials and Methods) score. The accuracies were calculated using k-fold cross-validation with *k* = 5, i.e. the data was split into training and testing sets 5 times, in such a way that each training example was in the test set once, and the prediction for each data point when it was in the test set was used. The results are shown for three classification algorithms, *ℓ*1-regularized logistic regression (LR), the random forest and a linear SVM. The regularized LR outperforms the random forest for many, though not all, functions. The average score over all functions for LR is 90.1% (versus 84.5% for the random forest), and 65 functions have a score greater than 90%, with 45 higher than 95%. The perfmormance of the SVM and the LR are similar, although they do differ significantly for some functions. The mean score of the SVM is slightly better, at 90.8% vs. 90.1%. This difference is not statistically significant (paired t test, p=0.71), and since LR is easier to interpret and much more computationally efficient, we decided to focus on it for the rest of the paper.

**Fig 1.**
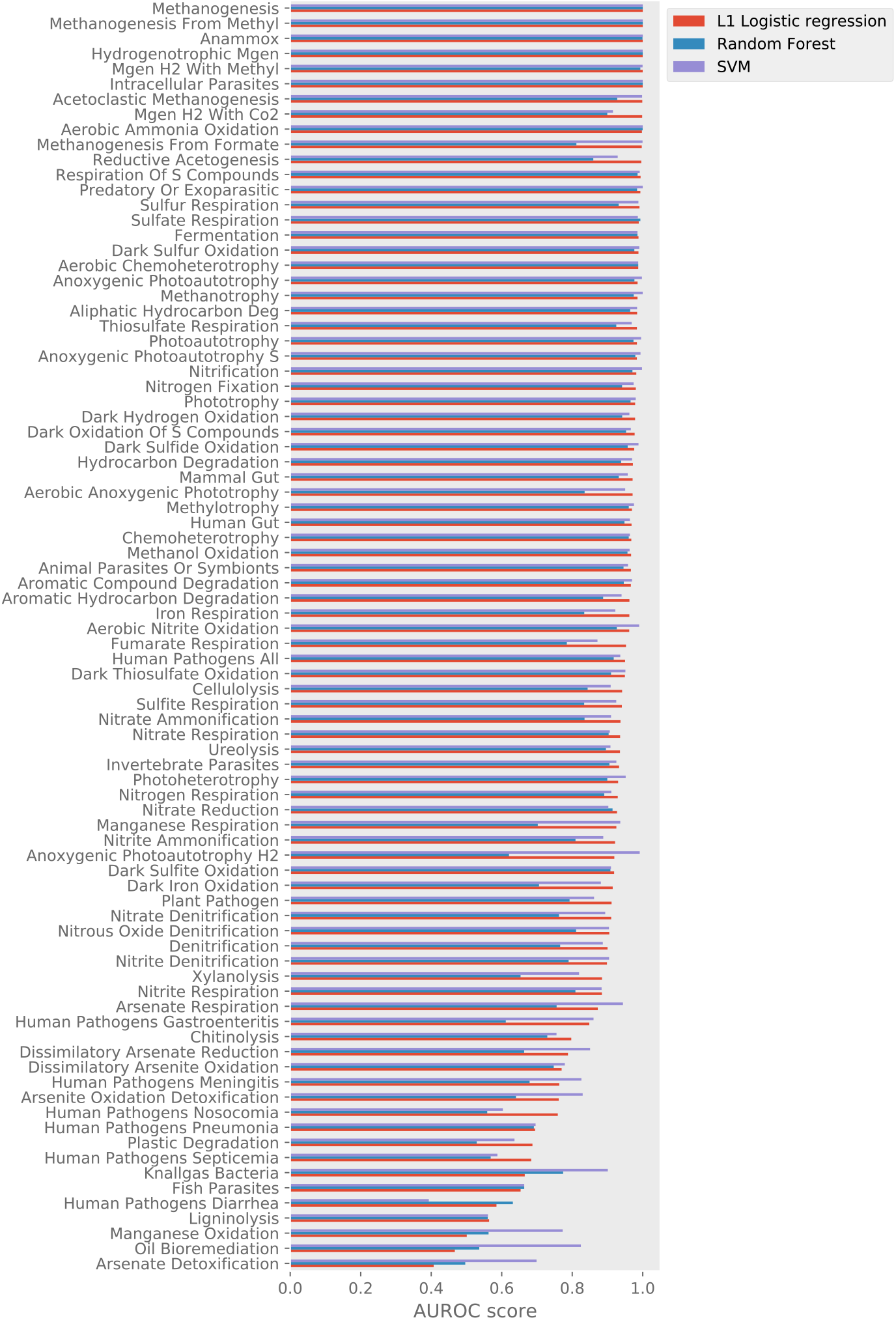
Overall performance of classification algorithms. The AUROC score on each classification task (each function) is shown for three classification algorithms: *ℓ*1-regularized logistic regression (LR), the random forest and a linear support vector machine (SVM). Functions are ordered by the LR score.

Additionally, we can compare the results we obtain using different gene ortholog schemes, that is KEGG orthologs vs. Pfam families. The results using the two schemes are rather similar, though there are some functions where one approach outperforms the other (Figure S1); this may reflect better coverage of the genes involved in the function in a particular scheme. On average, KO performs better, with a mean score of 90.1% versus 84.9% for Pfam (*p* < 0.001), and we therefore concentrate on the KO scheme for the remainder of the paper.

### Gene orthologs used by classifiers

Table 1 shows the KEGG orthologs with non-zero coefficients used by the logistic regression (LR) classifiers and their weights for some example functions. Due to the *ℓ*1-regularization, the number of non-zero coefficients is rather low. Three representative functions, all having classifiers with AUROC scores greater than 95%, are shown. Many of the KEGG orthologs picked out by the classifiers are genes known to be involved in these functions, as we might hope. In particular, consider the prediction of methanogens, a relatively easy task since it is known that methanogens must possess the *mcrA* gene, this being a necessary and sufficient condition for methanogenesis [45]. Indeed, subunits of this gene have the highest weight, and a total of only 9 genes are used by the classifier.

**Table 1.** Details of classifiers for specific functions. Tables showing all the nonzero weights in the logistic regression models trained on three functions from the FAPROTAX database. Note that there are 9647 KEGG orthologs used in our models, so the vast majority of weights are set to zero in these models.

| Sulfate respiration |  |  |
| --- | --- | --- |
| KO | Weight | Description |
| K03421 | 0.263 | methyl-coenzyme M reductase subunit C |
| K14109 | 0.226 | energy-converting hydrogenase A subunit R |
| K14094 | 0.156 | energy-converting hydrogenase A subunit C |
| K00401 | 0.131 | methyl-coenzyme M reductase beta subunit [EC:2.8.4.1] |
| K14097 | 0.093 | energy-converting hydrogenase A subunit F |
| K00440 | 0.091 | coenzyme F420 hydrogenase subunit alpha [EC:1.12.98.1] |
| K06862 | 0.045 | energy-converting hydrogenase B subunit Q |
| K16204 | 0.038 | seco-amyrin synthase [EC:5.4.99.52 5.4.99.54] |
| K11099 | 0.033 | small nuclear ribonucleoprotein G |
| K14098 | 0.027 | energy-converting hydrogenase A subunit G |
| K09613 | 0.026 | COP9 signalosome complex subunit 5 [EC:3.4.-.-] |
| K14093 | 0.022 | energy-converting hydrogenase A subunit B |
| K08074 | 0.013 | ADP-dependent glucokinase [EC:2.7.1.147] |
| K05181 | 0.013 | gamma-aminobutyric acid receptor subunit beta |
| K09493 | 0.013 | T-complex protein 1 subunit alpha |
| K06612 | 0.013 | alpha-N-acetyl-neuraminate alpha-2;8-sialyltransferase (sialyltransferase 8B) [EC:2.4.99.-] |
| K17278 | 0.011 | membrane-associated progesterone receptor component |
| K02938 | 0.003 | large subunit ribosomal protein L8e |
| K00442 | 0.003 | coenzyme F420 hydrogenase subunit delta |
| K14096 | 0.003 | energy-converting hydrogenase A subunit E |
| Methanogenesis |  |  |
| KO | Weight | Description |
| K03421 | 0.567 | methyl-coenzyme M reductase subunit C |
| K00400 | 0.276 | methyl coenzyme M reductase system; component A2 |
| K00579 | 0.160 | tetrahydromethanopterin S-methyltransferase subunit C [EC:2.1.1.86] |
| K00399 | 0.081 | methyl-coenzyme M reductase alpha subunit [EC:2.8.4.1] |
| K07463 | 0.023 | archaea-specific RecJ-like exonuclease |
| K17618 | 0.023 | ubiquitin-like domain-containing CTD phosphatase 1 [EC:3.1.3.16] |
| K00401 | 0.022 | methyl-coenzyme M reductase beta subunit [EC:2.8.4.1] |
| K09728 | 0.020 | uncharacterized protein |
| K09613 | 0.002 | COP9 signalosome complex subunit 5 [EC:3.4.-.-] |
| Hydrogenotrophic methanogenesis |  |  |
| KO | Weight | Description |
| K03421 | 0.231 | methyl-coenzyme M reductase subunit C |
| K14109 | 0.201 | energy-converting hydrogenase A subunit R |
| K00401 | 0.169 | methyl-coenzyme M reductase beta subunit [EC:2.8.4.1] |
| K14098 | 0.136 | energy-converting hydrogenase A subunit G |
| K14097 | 0.104 | energy-converting hydrogenase A subunit F |
| K14093 | 0.058 | energy-converting hydrogenase A subunit B |
| K06862 | 0.057 | energy-converting hydrogenase B subunit Q |
| K14094 | 0.049 | energy-converting hydrogenase A subunit C |
| K17278 | 0.043 | membrane-associated progesterone receptor component |
| K08074 | 0.042 | ADP-dependent glucokinase [EC:2.7.1.147] |
| K00442 | 0.032 | coenzyme F420 hydrogenase subunit delta |
| K09613 | 0.031 | COP9 signalosome complex subunit 5 [EC:3.4.-.-] |
| K09493 | 0.017 | T-complex protein 1 subunit alpha |
| K14099 | 0.009 | energy-converting hydrogenase A subunit H |
| K02938 | 0.005 | large subunit ribosomal protein L8e |
| K00399 | 0.001 | methyl-coenzyme M reductase alpha subunit [EC:2.8.4.1] |

Looking at some more complex traits, for example sulfate respiration (i.e. disimmilatory sulfate reduction to H_2_S), the model assigns a lot of weight to subunits of a quinone-modifying oxidoreductase, which is indeed associated with sulfur metabolism [46]. Interestingly, however, none of the genes picked out by the classifier are directly part of the metabolic pathway for this process as described in the KEGG module for dissimilatory sulfate reduction, see Figure 2. The situation is similar with hydrogenotrophic methanogenesis, with classification mostly determined by components of energy-converting hydrogenases which are not directly part of the autotrophic methanogenesis pathway, along with *mcr* genes indicating that the microbe is a methanogen.

**Fig 2.**
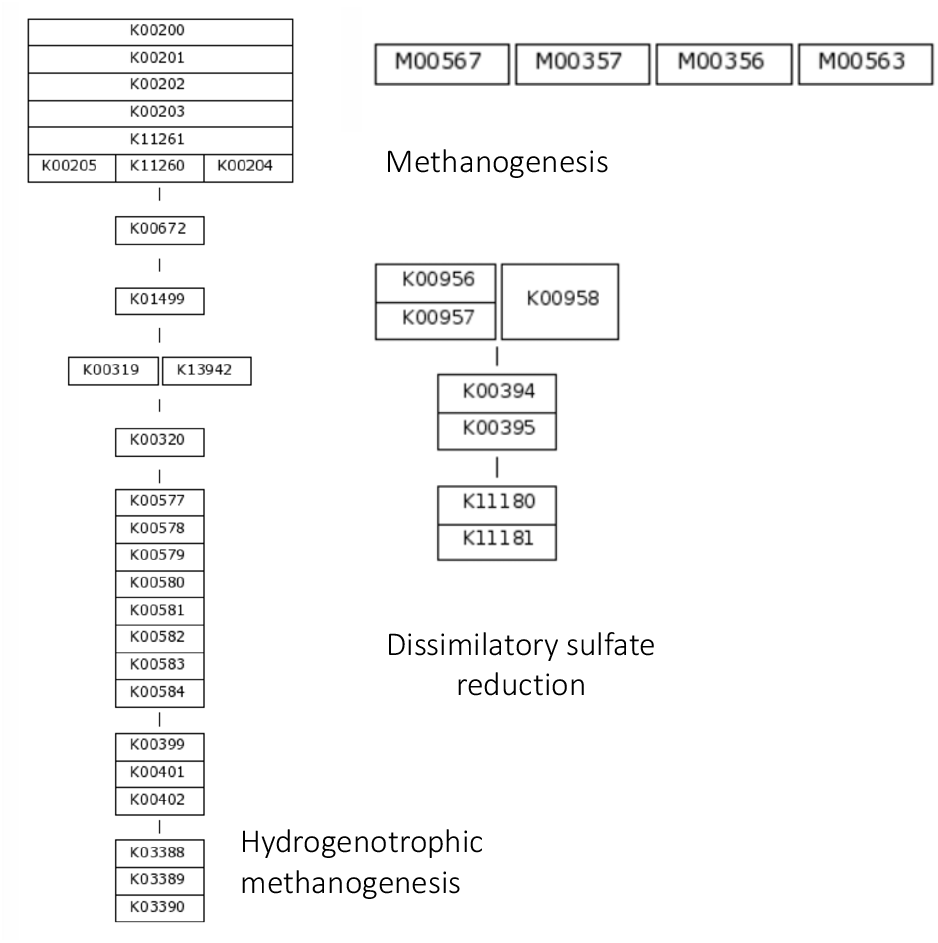
KEGG modules for some functions. Representations of the KEGG modules corresponding to the FAPROTAX functions shown in Table 1. Modules are organized into ‘blocks’ of orthologs, typically indicating a protein complex. Orthologs positioned next to each other are ‘options’, i.e. that section of the module is present if any of the adjacent blocks are present.

Figure 3 shows a scatter plot of AUROC score (i.e. classifier performance) against the number of orthologs used to make the prediction. It can be seen that there is a correlation between these two variables, with some highly accurate classifiers built out of a large number of genes. However, there is also a noticable cluster of functions with high accuracy achieved with only a few genes (less than 100). These functions may be particularly interesting, as it is more likely that these small groups of orthologs are causally associated with the function, rather than just being genes which typically occur in parts of the phylogenetic tree which have the function and may or may not have any direct relation to it. This issue is explored further in the section on performance across taxa, below. Also, note that most of the functions that perform poorly, which typically use very few genes to classify, have very low support in the training data in terms of number of positive examples.

**Fig 3.**
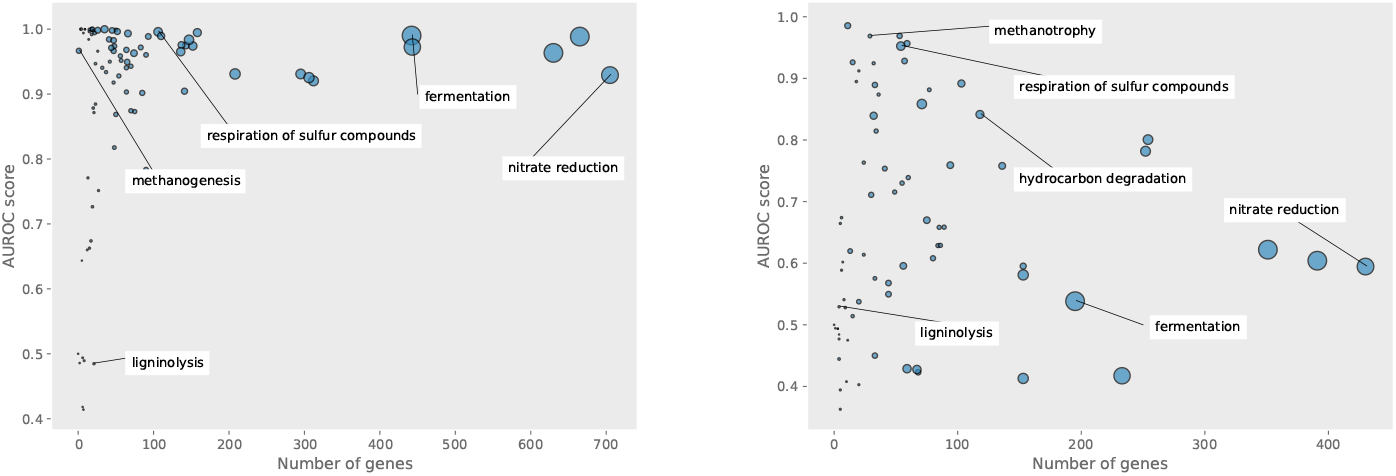
Scatterplots showing the AUROC score of the different classifiers plotted against the number of gene orthologs the classifier uses to make its predictions. Point size is proportional to the number of positive examples in the training set. Left: in the standard case. Right: in the cross-taxa case.

Figure S2 shows an ordination plot of all the species in the training and test sets using their KEGG ortholog copy numbers. That is, a dimensionality reduction algorithm, here stochastic neighbour embedding, has been applied to visualise variation in all KO copy numbers in two dimensions. Points are colored by whether they are true positive, true negative, false positive or false negatives under a particular classification task, here for sulfate respiration. It can be seen that the species performing this function do tend to cluster together into a few groups in the KO space, allowing our algorithm to classify them mostly correctly.

### Comparison to KEGG modules

It is instructive to compare the performance of our classifiers to the use of KEGG modules, where an equivalent module exists for that function, i.e. compare the performance to a ‘classifier’ where an organism is judged capable of a function if it has a complete KEGG module for that function. Table 2 shows the results of this comparison for three FAPROTAX functions with corresponding KEGG modules. Note that the KEGG module method does not require training, so the metrics are over the entire NCBI dataset, whereas for the classifier they are only for the held-out test set. Also, the former method gives only presence/absence of a function rather than a probability, so the AUROC score cannot be calculated, so we use alternative metrics based on classification: the *F*_1_ score and the confusion matrix.

**Table 2.** Comparison of classifiers to KEGG modules. Table showing the performance of using KEGG module presence/absence against LR classifiers for some functions where equivalent KEGG modules exist. Since the KEGG module approach does not give a probability, the AUROC score cannot be used, so the F1 score and confusion matrices are compared.

| module | KEGG modules |  | Classifier |  |
| --- | --- | --- | --- | --- |
|  | F1 | confusion matrix | F1 | confusion matrix |
| sulfate respiration | 0.84 | $\begin{pmatrix} 9197 & 53 \\ 6 & 151 \end{pmatrix}$ | 0.99 | $\begin{pmatrix} 2313 & 0 \\ 1 & 38 \end{pmatrix}$ |
| nitrate respiration | 0.14 | $\begin{pmatrix} 6821 & 2162 \\ 224 & 200 \end{pmatrix}$ | 0.622 | $\begin{pmatrix} 2237 & 9 \\ 54 & 52 \end{pmatrix}$ |
| hydrogenotrophic methanogenesis | 0.756 | $\begin{pmatrix} 9281 & 43 \\ 6 & 77 \end{pmatrix}$ | 0.923 | $\begin{pmatrix} 2331 & 0 \\ 3 & 18 \end{pmatrix}$ |

It can be seen that the LR classifier does significantly better than KEGG modules in assigning these functions as they appear in the FAPROTAX database. This suggests that having the enzymes or proteins described in the KEGG module for a function is not in fact a necessary or sufficient condition for actually performing that function, and that other genes are more predictive. However, it is possible that the discrepancy is due instead to inaccuracy in the FAPROTAX database, e.g. species which do perform the functions being missed from the database and therefore getting flagged as false positives with the KEGG method. More work would be needed to fully exclude this possibility.

### Performance across taxa

As mentioned above, it is not clear how much the genes being used by the classifiers are related to the functions being predicted or are reflecting taxonomic relations among phylogenetically similar organisms that perform the same or similar functions. To assess this, we analysed how predicted functions are distributed across a taxonomic orders. We found that this distribution varies with function, where some functions are taxonomically clustered while others are not (Figure 4) 4(as also seen in other studies [2]).

**Fig 4.**
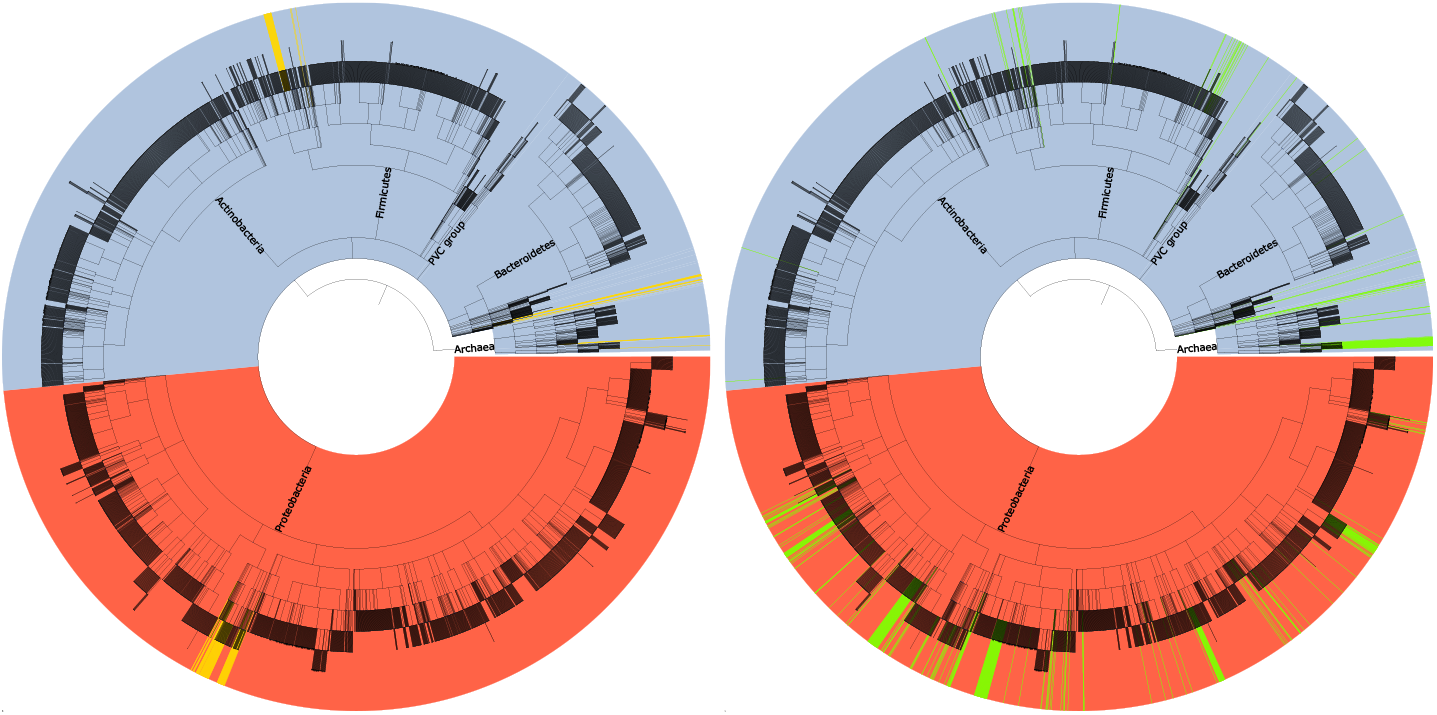
Taxonomic distribution of metabolic traits. Taxonomic trees of all prokaryotic NCBI species with full genomes. For training the cross-taxa verison of the classifier, only the Proteobacteria (red section of the tree) were used, and the models were tested on the rest of the tree. Species capable of a) sulfate respiration and b) nitrate respiration are highlighted on the trees.

To see if we can develop classifiers that are less affected by taxonomic signals, we trained the models on organisms from a subset of taxa and tested its performance on another, unrelated taxa. If a classifier can predict phenotype based on genes in a distantly-related, unseen set of organisms, it is likely the genes it is using have a real association with the function. In particular, we tried training our logistic regression models on the Proteobacteria, a large phylum of bacteria, and testing on the rest of the taxonomy. Some functions did not have significant numbers of species in each of these sets; we used only functions with at least 5 species in the training set and 5 in the test, leaving 59 functions out of 84.

As might be expected, the resulting classifiers from this approach performed significantly worse in this case, compared to being trained on a random selection of species from throughout the prokaryotic part of the tree of life, see Figure 3b. However, for a significant number of functions the performance of the classsifier is still fairly good, indicating an ability to make predictions which are generalizable to significantly different unseen groups of organisms. 19 functions have an AUROC score greater than 80%, and 9 greater than 90%.

Figure 3b shows a scatter plot of classifier complexity against performance, as in Figure 3a. Notable is that the group of classifiers achieving high accuracy while using a lot of genes is gone: functions such as fermentation and nitrate reduction, which were in this group of classifiers, are now much less accurate. Classifiers which work well in the cross-taxa case all use a relatively small number of genes, less than 150 or so. This suggests that the classifiers using a large number of genes to make predictions in the randomized case may have been using a range of genes found in different closely-related clusters of organisms which all have the target trait, but which may not have a causal relationship with the function.

### Prediction of MAG phenotypes

A possible major benefit of developing machine learning approaches to predicting function from genomes is that the resulting classifiers can be applied to novel genomes predicted from meta genomes. We therefore used the classifiers trained above to classify metagenomically assembled genomes (MAGs) from a few different environments. These were laboratory anaerobic digesters, the ocean (from the Tara oceans project [5]), and MAGs from a groundwater aquifer assigned to be members of the so-called ‘canditate phyla radioation’ (CPR) [8]. The CPR is a set of bacterial lineages discovered from metagenomic studies consisting of a very large number of proposed novel phyla. These organisms have very small genomes, and may typically live in symbiosis with other organisms [47].

To perform the functional assigments, we used the *ℓ*1-regularized LR classifier described above, with a random train-test splitting and the regularization parameter *C* = 0.05, trained using KEGG orthologs on the full NCBI genomes. Figure 5 shows the predicted functions by this classifier for the MAGs assembled from anaerobic digesters and from the global oceans. There are noticeable differences, such as more AD MAGs having fermentation and sulfate-metabolism-related functions and fewer having aerobic chemoheterophy.

**Fig 5.**
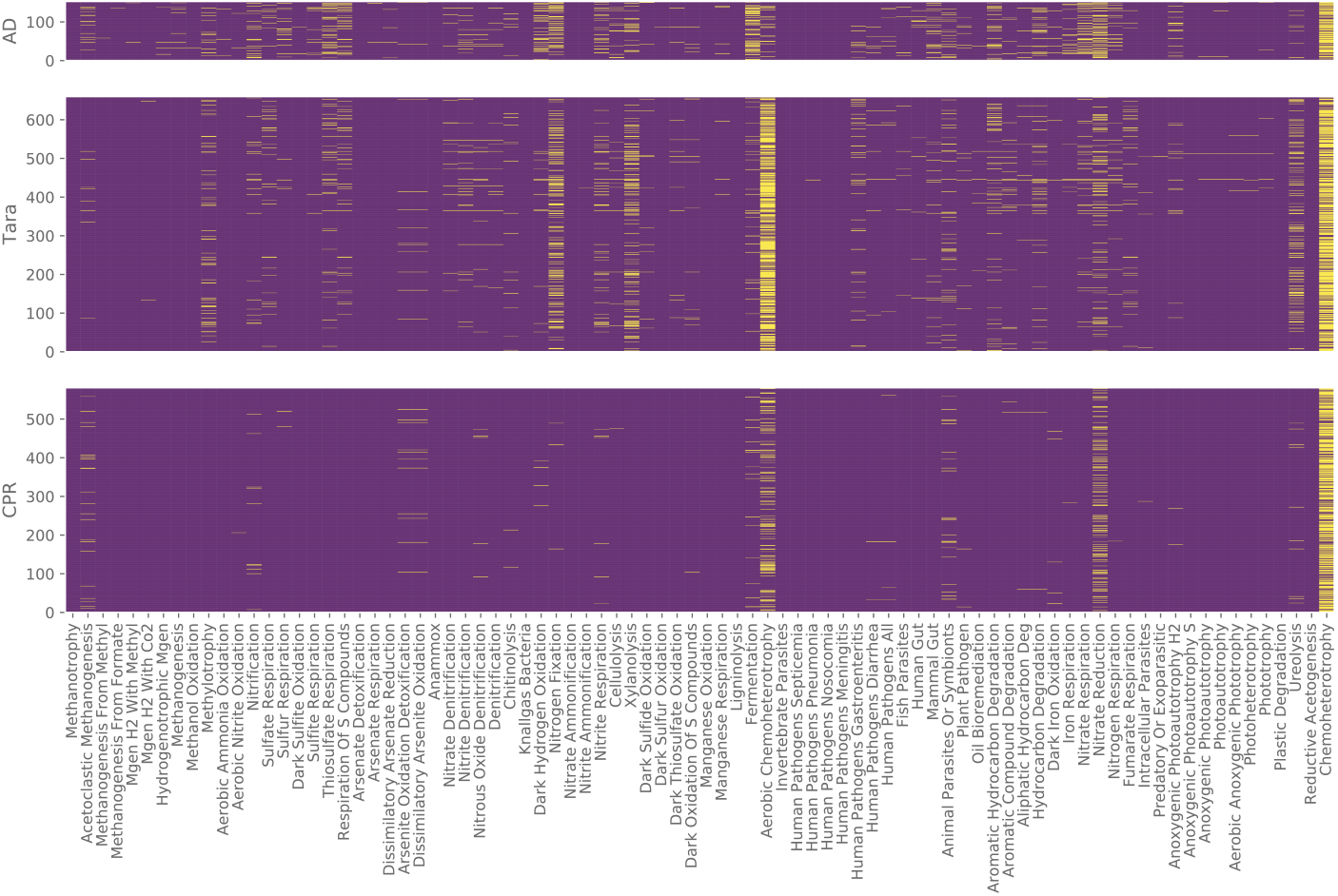
Heatmap of presence/absence of fucntions in MAGs. Results of running the set of LR classsifiers trained on NCBI genomes on MAGs assembled from three environments: laboratory anaerobic digesters, the ocean and ‘candidate phyla radiation’ (CPR) organisms from an aquifer system.

To make these differences clearer, Figure S3 is a bar chart showing the proportion of MAGs from the different environments having a function, for some of the most common functions. For many functions, the differences are very significant.

These differences in function might well be expected between these environents. For example, fermentation is very important in the AD process, and aerobic chemoheterotrophy obviously is not as the environment is aerobic. This indicates that the method is capable of producing useful information about MAGs. The results for the CPR MAGs indicate that these organisms possess significantly fewer functions than those from the other two environments, as would be expected from their very small genome sizes. A few functions do however have significant incidence in this group. Apart from ‘chemoheterotrophy’ and ‘aerobic chemoheterotrophy’, which are very broad categories encompassing a large proportion of all organisms, a few functions associated with nitrogen metabolism, especially nitrate reduction, are noticeably present in the group. That the CPR are involved in nitrate reduction was recently proposed in Danczak et al. [47].

Some of the functional assignments seem strange, for example organisms being classified as acetoclastic or hydrogenotrophic methanogens but not as methanogens, including a significant proportion of CPR organisms (about 3%). Looking at the gene orthologs present in these organisms and their taxonomic assignments sheds some light on what is going on here, see Table 3. For example, some of the acetoclastic methanogens which are misclassified as not being methanogens are missing the *mcrA* ortholog K00399, presumably because the MAGs are incomplete and this gene has been missed. Another example is an organism classified as being a hydrogenotrophic methanogen but not a methanogen. This MAG appears to be similar to the genome of the bacterium *Caldisericum exile*, which is not a methanogen and does not possess *mcrA* (it is an anaerobic, thermophilic bacterium which respires by thiosulfate reduction ??). However, it does possess genes for subunits of the energy-converting hydrogenase A, which is indicative of hydrogenotrophic methanogenesis (see Table 1). Therefore, these discrepancies may be the result either of incomplete MAGs, or of combinations of genes which are rare or unseen in the training set.

**Table 3.** Key genes and predicted functions for MAGs predicted to be methanogens. Gene copy numbers for the *mcrA* methanogenesis gene and the energy-converting hyrdogenase A, along with functional predictions, for AD MAGs predicted to be methanogenic by our algorithm.

| Species | mcrA | echA | methanogenesis | hydrogenotrophic | acetoclastic |
| --- | --- | --- | --- | --- | --- |
| Caldisericum exile | 0 | 1 | 0 | 1 | 0 |
| Methanomassiliicoccus luminyensis | 1 | 0 | 1 | 1 | 0 |
| Methanosaeta concilii | 2 | 0 | 1 | 1 | 1 |
| Methanosaeta concilii | 0 | 0 | 1 | 0 | 1 |
| Methanosaeta harundinacea | 1 | 0 | 1 | 0 | 0 |
| Methanoregula formicica | 1 | 0 | 1 | 0 | 0 |
| Methanoregula formicica | 1 | 0 | 1 | 0 | 0 |
| Methanolinea tarda | 1 | 0 | 1 | 0 | 0 |

For the Tara dataset, metadata for different samples was available, including depth, temperature and salinity, among other measurements. Combining the coverages of the different MAGs across these samples with the functional assignments of the MAGs, we can calculate the proportion of a given function in different sample groups. Figure 6 shows the mean of this metric for all the functions for samples from different oceans. The functional differences do not seem to be very significant between oceans, with a few exceptions, notably an abundance of nitrate reduction in the Southern Ocean, and a strong negative correlation between temperature and fermenter abundance (Pearson correlation *r* = –0.43, *p* < 0.001). Figure 7.

**Fig 6.**
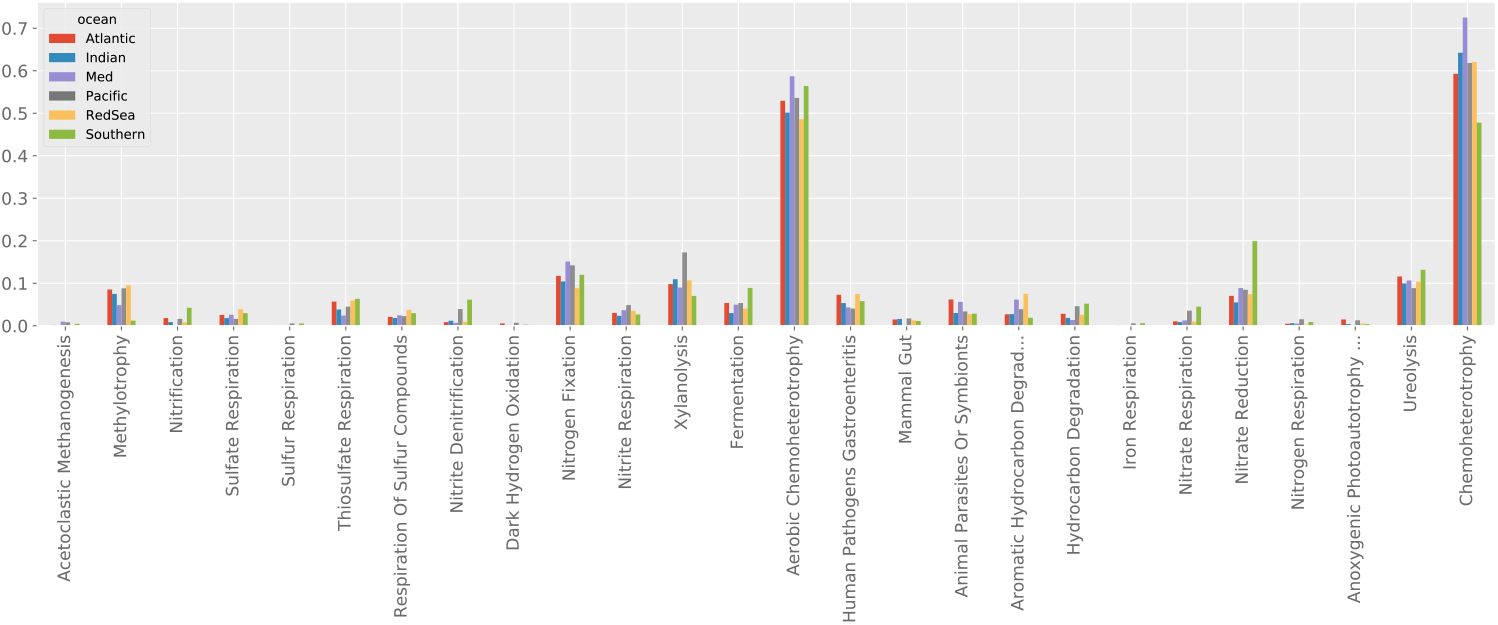
Mean abundance of microbes performing functions by ocean. Average of the proportion of total coverage associated with microbes performing a function over samples from each of the oceans.

**Fig 7.**
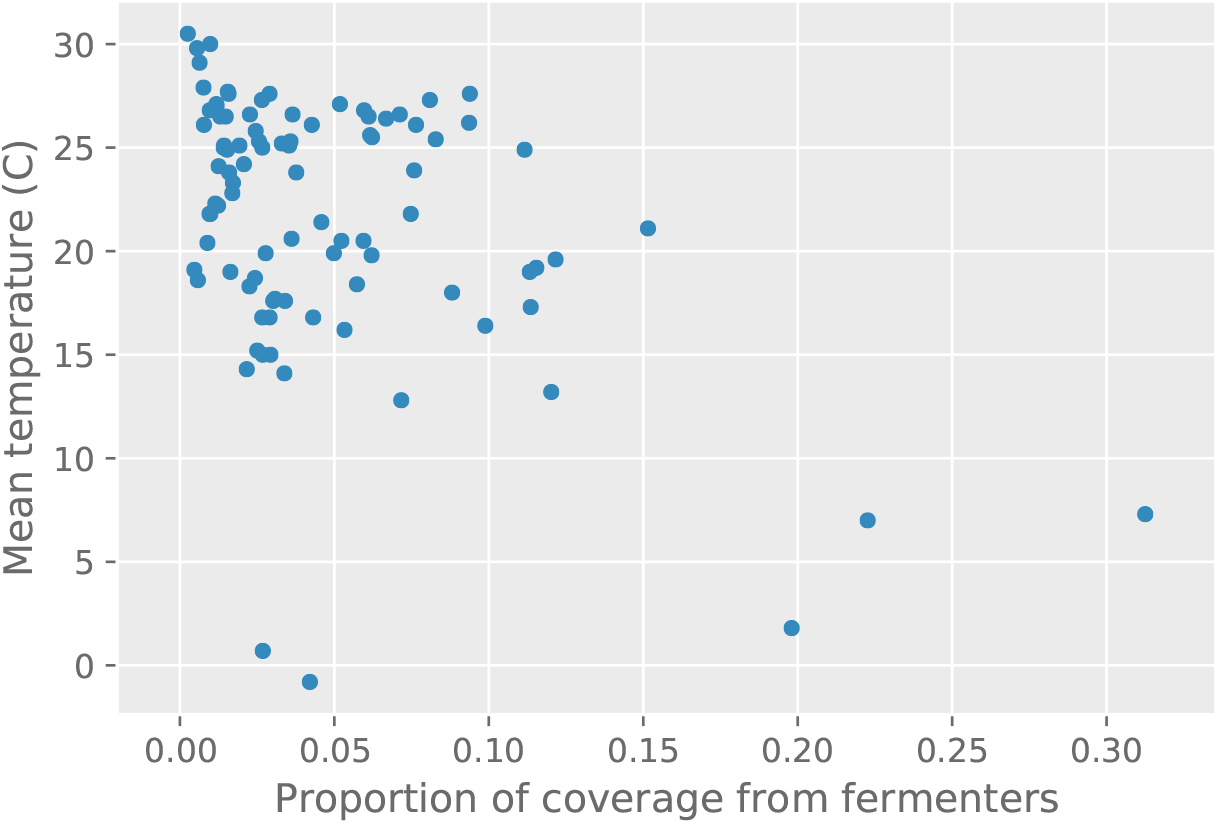
Association of temperature and fermenter abundance. Scatterplot of proportion of total coverage associated with microbes performing fermentation versus mean temperature of a sample. Pearson *r* = –0.43, *p* < 0.001.

## Discussion

We have demonstrated a machine learning method for inferring phenotypes from genomes. This method uses gene orthology and copy numbers as its features and is trained using over 9000 genomes and their known functional phenotypes. While the accuracy of the predictions vary significantly over different phenotypes, a significant proportion of the functions we tested achieved very good classification accuracy, with AUROC scores greater than 90%. Of the machine learning algorithms we tested, we found that *ℓ*1-regularized logistic regression gave the best combination of accuracy, computational efficiency and interpretability. The results did not depend very strongly on whether KEGG orthologs or Pfam domains were used to characterise genes in the genomes, although the KEGG scheme performed slightly better on average over the functions we considered here. The logistic regression models we generate can be inspected, and the genes most associated with a given phenotype in the model enumerated, which allows for validation of the models by comparison to what is known about the function of these genes. This raises the potential for this method to discover new associations between orthologous groups and phenotypes. For example, we found here that the presence of subunits of the energy-converting hydrogenase A are more predictive of an organism’s performing hydrogenotrophic methanogensis than the genes directly involved in the process as described in the KEGG functional module for it. For nitrate reduction, in addition to expected genes such as nitrate reductase, there are multiple KEGG orthologs listed as ‘uncharacterised protein’ which are highly predictive of this function.

To check the robustness of the models we generated, we tried training the models on one section of the microbial taxonomic groups, the Proteobacteria, and testing its accuracy on organisms from the rest of the tree, which it had not encountered at all in training. This did significantly reduce model accuracy for many phenotypes. This is to be expected, as the training and test sets in this case are so different. However, some of the functions still achieved good accuracy. This would suggest that the logistic regression model is identifying genes functionally involved with the phenotype in the training stage, such that their presence even in distantly related organisms is indicative of the presence of the phenotype. Phenotype predictions that had good accuracy under this scheme tended to produce models involving only a few genes (i.e. only a few genes had nonzero weights in the logistic regression model), less than 100, supporting the idea that these models are picking out genes directly involved with the phenotype.

As with any machine learning approach, the presented study is limited by the accuracy and availability of the data on which models can be trained. We consider this to be the key bottleneck in the genotype-phenotype prediction problem and expect future improvements to come from better data curation and collection (on existing organsisms), rather than development of new improved classification algorithms. To this end, extended versions of databases such as FAPROTAX, and listing detailed phenotypic features of cultured organisms would provide a highly valuable resource for machine learning approaches. Such data would allow development of finer grained and more accurate classifiers, which can then be applied to unknown genomes and MAGs. The results of such applications will also be improved by better MAG assembly. In particular, we note the key limitation of classifiers on MAGs being the accuracy of their gene content assignments. Increasing this accuracy will also improve the accuracy of their machine learning based functional assignment. As these limitations are addressed, we expect classifiers to become a key tool in gaining insights into the functional capabilities both of microbiomes as a whole and the individual species making up those communitites, even when those species cannot be cultivated.

## Supporting information

**Fig S1.**
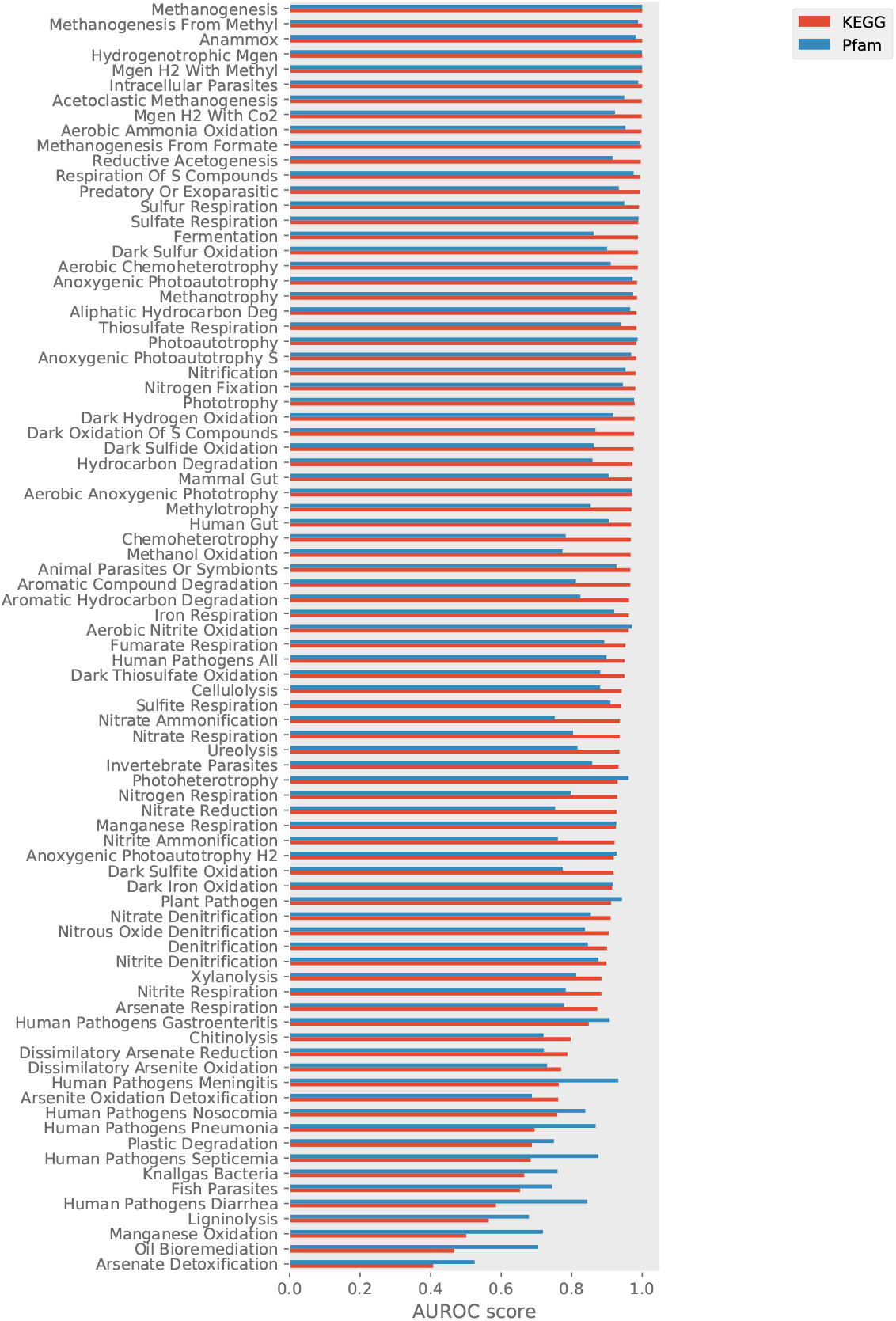
Performance of classification algorithms using different ortholog schemes. The AUROC score on each classification task (each function) is shown for algorithms trained on two representations of genomes in terms of orthologous groups of genes, the KEGG orthology (KO) and Pfam protein families.

**Fig S2.**
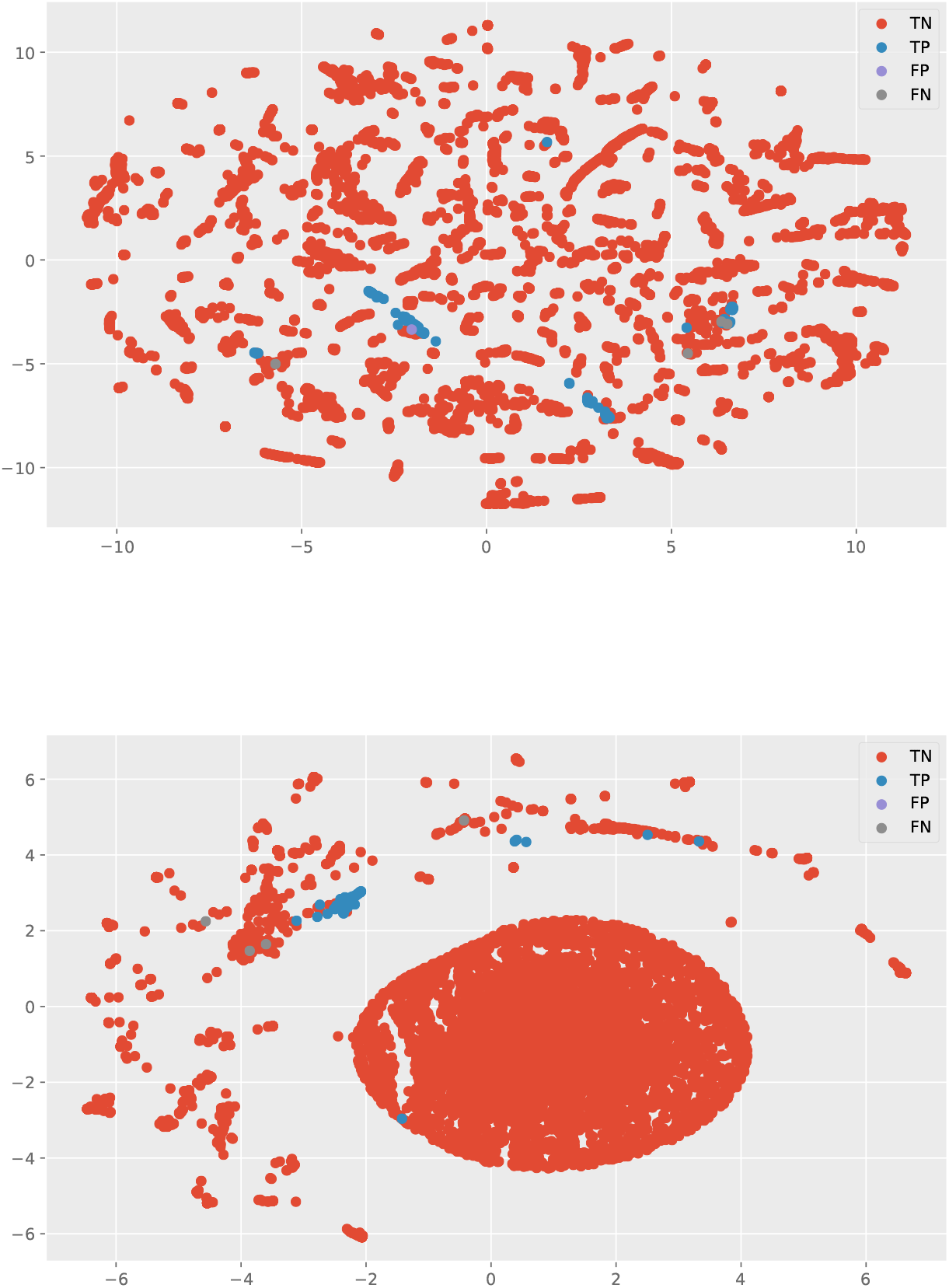
Stochastic neighbour embedding of all species in KEGG ortholog space. Ordination in two dimensions of all the species in our training dataset. By coloring by classification group (true positive, true negative, false positive, false negative) for a particular function, here sulfate respiration, we can graphically visualise the behaviour of our classifier. Top: embedding performed over all KEGG orthologs. Bottom: embedding performed only over KOs relevant to the function according to the classifier.

**Fig S3.**
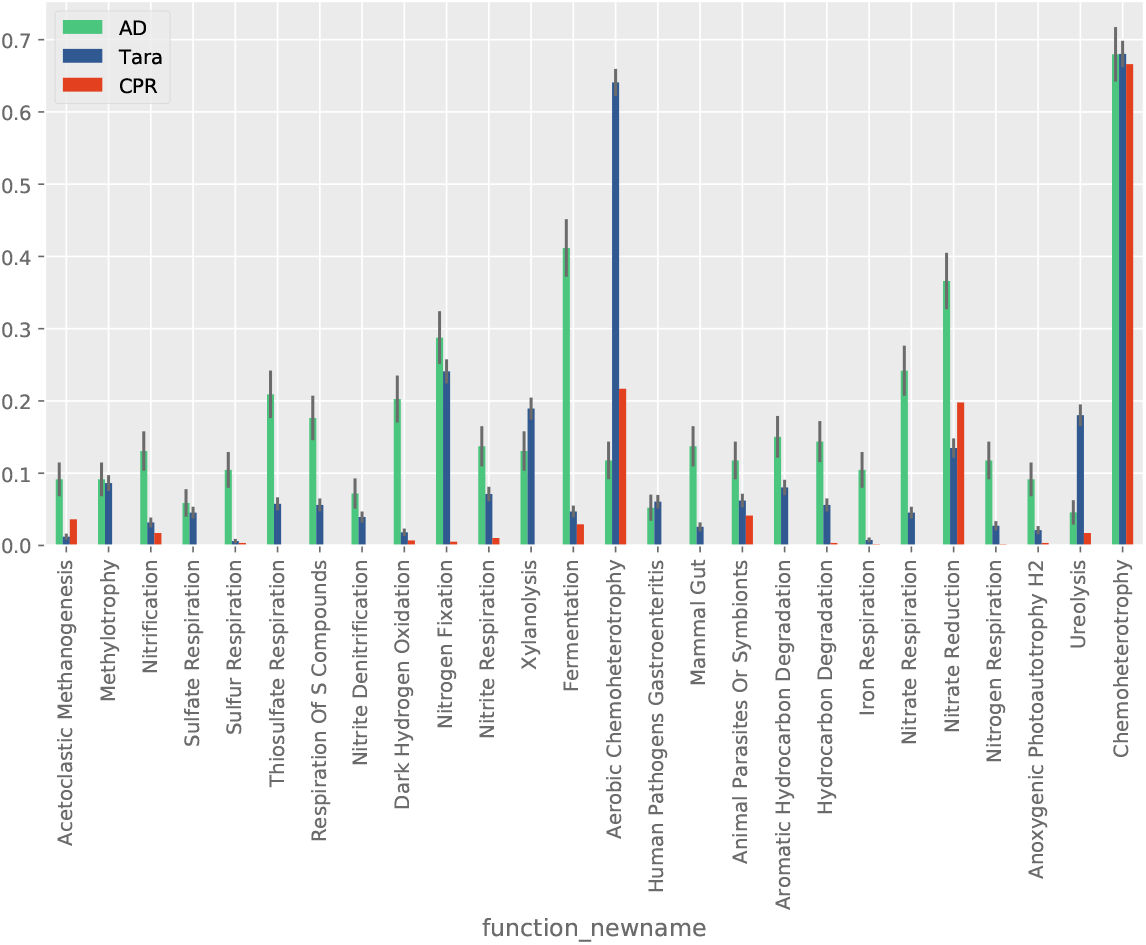
Overall comparison of AD, Tara and CPR MAGs. Proportions of MAGS from the environments having a function, for some of the most common functions.

